# Micrococcin cysteine-to-thiazole conversion through transient interactions between a scaffolding protein and two modification enzymes

**DOI:** 10.1101/2023.10.23.563616

**Authors:** Diana G Calvopina-Chavez, Devan M Bursey, Yi-Jie Tseng, Leena M Patil, Kathryn D Bewley, Philip R Bennallack, Josh M McPhie, Kimberly B Wagstaff, Anisha Daley, Susan M Miller, James D Moody, John C Price, Joel S Griffitts

## Abstract

Ribosomally synthesized and post-translationally modified peptides (RiPPs) are a broad group of compounds mediating microbial competition in nature. Azole/azoline heterocycle formation in the peptide backbone is a key step in the biosynthesis of many RiPPs. Heterocycle formation in RiPP precursors is often carried out by a scaffold protein, an ATP-dependent cyclodehydratase, and an FMN-dependent dehydrogenase. It has generally been assumed that the orchestration of these modifications is carried out by a stable complex including the scaffold, cyclodehydratase and dehydrogenase. The antimicrobial RiPP micrococcin begins as a precursor peptide (TclE) with a 35-amino acid N-terminal leader and a 14-amino acid C-terminal core containing six Cys residues that are converted to thiazoles. The putative scaffold protein (TclI) presumably presents the TclE substrate to a cyclodehydratase (TclJ) and a dehydrogenase (TclN) to accomplish the two-step installation of the six thiazoles. In this study, we identify a minimal TclE leader region required for thiazole formation, we demonstrate complex formation between TclI, TclJ and TclN, and further define regions of these proteins required for complex formation. Our results point to a mechanism of thiazole installation in which TclI associates with the two enzymes in a mutually exclusive fashion, such that each enzyme competes for access to the peptide substrate in a dynamic equilibrium, thus ensuring complete modification of each Cys residue in the TclE core.

**IMPORTANCE:** Thiopeptides are a family of antimicrobial peptides characterized for having sulfur-containing heterocycles and for being highly post-translationally modified. Numerous thiopeptides have been identified; almost all of which inhibit protein synthesis in gram-positive bacteria. These intrinsic antimicrobial properties make thiopeptides promising candidates for the development of new antibiotics. The thiopeptide micrococcin is synthesized by the ribosome and undergoes several post-translational modifications (PTMs) to acquire its bioactivity. In this study, we identify key interactions within the enzymatic complex that carries out cysteine to thiazole conversion in the biosynthesis of micrococcin.

## INTRODUCTION

Ribosomally synthesized and post-translationally modified peptides (RiPPs) are natural products produced by many bacteria that exhibit diverse biological activities including antimicrobial functions (1–6). RiPP biosynthesis starts with the ribosomal translation of a precursor peptide that is then heavily modified by multiple enzymes. The precursor peptide consists of an N-terminal leader sequence, also known as the recognition sequence, responsible for recruiting enzymes that carry out post-translational modifications (PTMs) (7–9), and a C-terminal core peptide sequence where PTMs occur (10–12). In most cases, RiPP biosynthetic gene clusters encode an E1-ubiquitin activating-like (E1-like) protein that has been implicated in leader peptide binding. This E1 homolog contains a RiPP recognition element (RRE) that adopts a highly conserved winged helix-turn-helix (wHTH) structure with three α-helices and a three-stranded β-sheet. RiPP leader peptides bind to RRE domains by interacting at the interface of the 3α/3β fold acting as a fourth β strand (13–15). After proper substrate recognition, numerous possible modifications take place on the core peptide culminating with the proteolytic removal of the leader from the core yielding a mature RiPP (7, 8).

Thiazole/oxazole-modified microcins (TOMMs) are a class of RiPPs that feature thiazol(in)e and oxazol(in)e heterocycles resulting from intramolecular reactions of cysteine, serine or threonine residues in the precursor peptide (16). Thiazole/oxazole biosynthesis is a two-step process in which an ATP-dependent cyclodehydratase (member of the YcaO superfamily) yields thiazoline/oxazoline heterocycles that are then oxidized into azoles by an FMN-dependent dehydrogenase. In addition to the cyclodehydratase and optional dehydrogenase, TOMM clusters encode proteins that facilitate coupling of the precursor peptide with these enzymes, but the different TOMM systems are highly variable in this respect. Most include an E1-like scaffold protein (mentioned above) and/or a second type of protein-protein interaction domain annotated as “Ocin-ThiF-like”. Either or both of these may contain an RRE, and the E1-like domain is often fused to the cyclodehydratase (17–23). This structural variability in TOMM complexes is illustrated in **Fig. 1**, which depicts four examples for which studies on the architecture of these complexes have been carried out. During biosynthesis of the cyanobactin trunkamide, the enzyme TruD catalyzes formation of azoline heterocycles. The crystal structure of TruD shows a fused cyclodehydratase with an NTD that contains an E1-like domain with an RRE, while the CTD comprises a YcaO domain responsible for catalyzing heterocycle formation (24, 25). The biosynthetic gene cluster for the bacteriocin heterocycloanthracin (HCA) contains a single copy of an Ocin-ThiF-like protein (HcaF), a fused E1-YcaO cyclodehydratase (HcaD), and a dehydrogenase (HcaB). In this system, the Ocin-ThiF-like protein interacts in a 1:1 ratio with the E1-like domain on the fused cyclodehydratase to yield azoline heterocycles. These azoline rings are then oxidized into azoles by a dedicated dehydrogenase, however, studies to characterize protein interactions with this enzyme have been unsuccessful (26, 27). For Microcin B17, the E1-like scaffold protein (McbB) interacts with a discrete cyclodehydratase (McbD) and a dehydrogenase (McbC) in a higher order octameric complex with a ratio of 4:2:2 (22, 28). In a more complicated system, the biosynthetic gene cluster for the thiopeptide sulfomycin (Sul) encodes multiple copies of RREs (SulB, SulF), E1/Ocin-ThiF-like proteins (SulB, SulC, SulE, SulF), cyclodehydratases (SulC, SulD) and dehydrogenases (SulF, SulG). Combinations of these proteins form three complexes (SulBC, SulEFG, SulDEFG) that achieve Cys, Thr and Ser conversion into their corresponding thiazole, methyloxazole, and oxazole (23, 29). All these TOMM systems share certain biochemical features across a vast evolutionary distance, but they vary in their intersubunit architectures.

**Figure 1.**
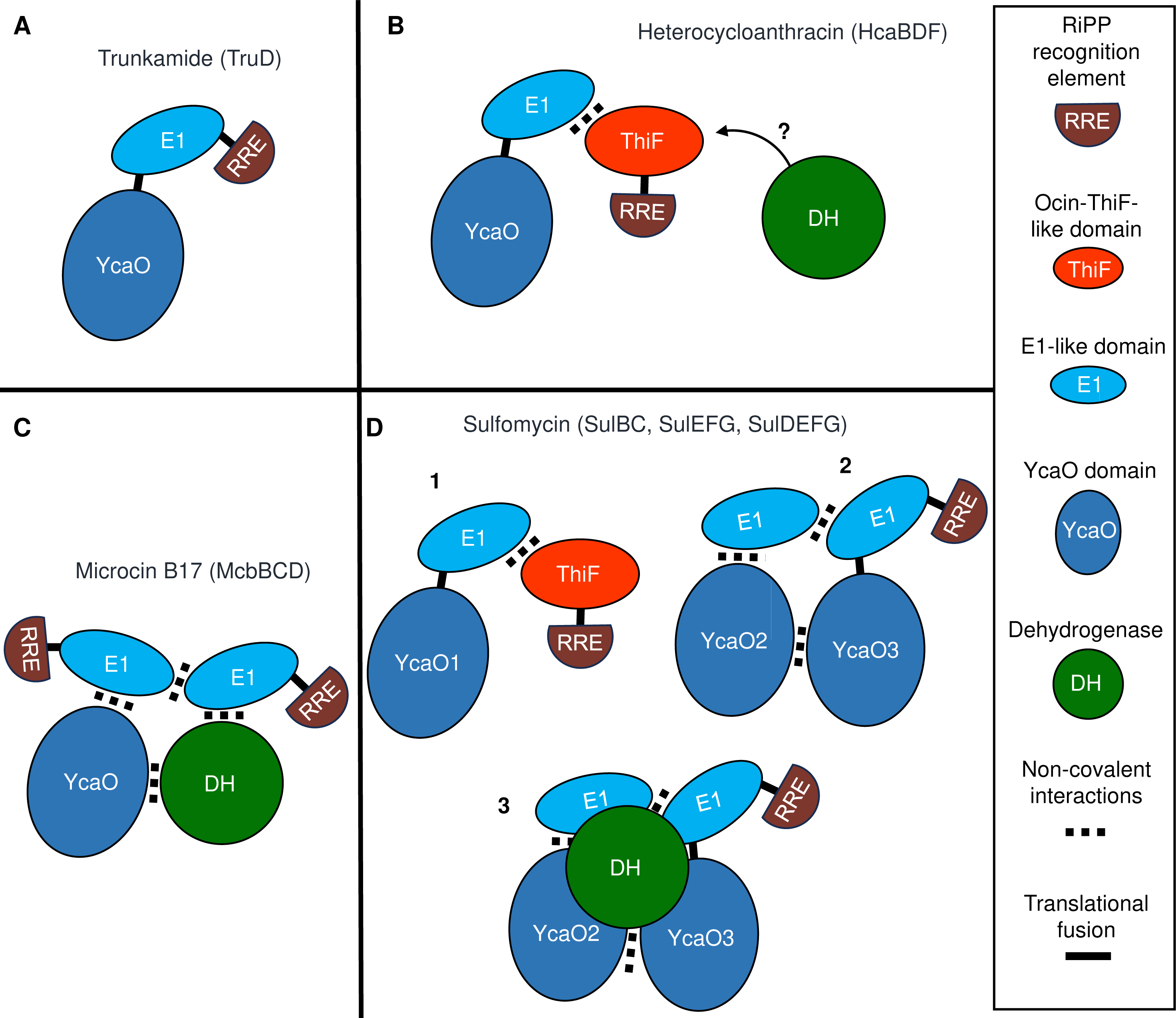
Schematic models of TOMM complexes from four RiPP biosynthetic pathways for which the structural architecture has been elucidated. Color codes for each protein are described on the box on the right. The cyclodehydratase enzyme from the YcaO superfamily is labeled YcaO. Black lines represent fused proteins, and dotted rectangles denote non-covalent interactions. A. Cartoon representation of the cyclodehydratase form the Trunkamide biosynthetic pathway (PDB: 4BS9) (18). B. Interactions between TOMM proteins from the microcin Heterocycloanthracin. The question mark denotes that the dehydrogenase is part of this pathway but attempts to characterize their protein interactions have not been successful (26). C. Cartoon diagram based on the crystal structure of Microcin B17 (PDB: 6GRI) (22) D. Paradigm of interactions between TOMM proteins in the biosynthesis of the thiopeptide sulfomycin. Heterocycle formation in this system is mediated by three complexes labeled 1, 2, and 3 (23).

Micrococcin is a thiopeptide produced by several Gram-positive bacteria, including *Bacillus cereus* and *Macrococcus caseolyticus* (30–32). Its biosynthesis involves several PTMs, including thiazole formation, C-terminal decarboxylation, dehydroamino acid formation, and the creation of a pyridine-anchored macrocycle (33). The gene cluster responsible for micrococcin production in *M. caseolyticus* (**Fig. 2A**) is located on a plasmid and consists of 12 *tcl* genes, which is simpler than the 24-gene cluster found in *Bacillus cereus* (34, 35). Out of these 12 *tcl* genes, 8 are essential for micrococcin production (31, 33). The roles of these genes are illustrated in **Fig. 2B**. The precursor peptide for micrococcin, TclE, has an N-terminal leader of 35 amino acids and a C-terminal core of 14 amino acids. Its biosynthesis begins with the conversion of all six cysteine residues in the core to thiazoles (31). Thiazole installation is required for all subsequent modifications (**Fig. 2B**). The work presented here focuses on the thiazole installation step in micrococcin biosynthesis. Each thiazole conversion is a two-step process requiring three proteins: a putative scaffold (TclI), a cyclodehydratase (TclJ), and a dehydrogenase (TclN) (**Fig. 2C**). We have previously shown that in the absence of TclI, no formation of thiazolines or thiazoles occurs, suggesting that this putative scaffold protein is essential for cys-to-thiazole conversion. When the TclJ cyclodehydratase is absent, there are no detectable thiazolines or thiazoles. In the absence of the TclN dehydrogenase, all six thiazoline heterocycles accumulate, suggesting that each Cys residue does not require complete modification before the next one is processed (31).

**Figure 2.**
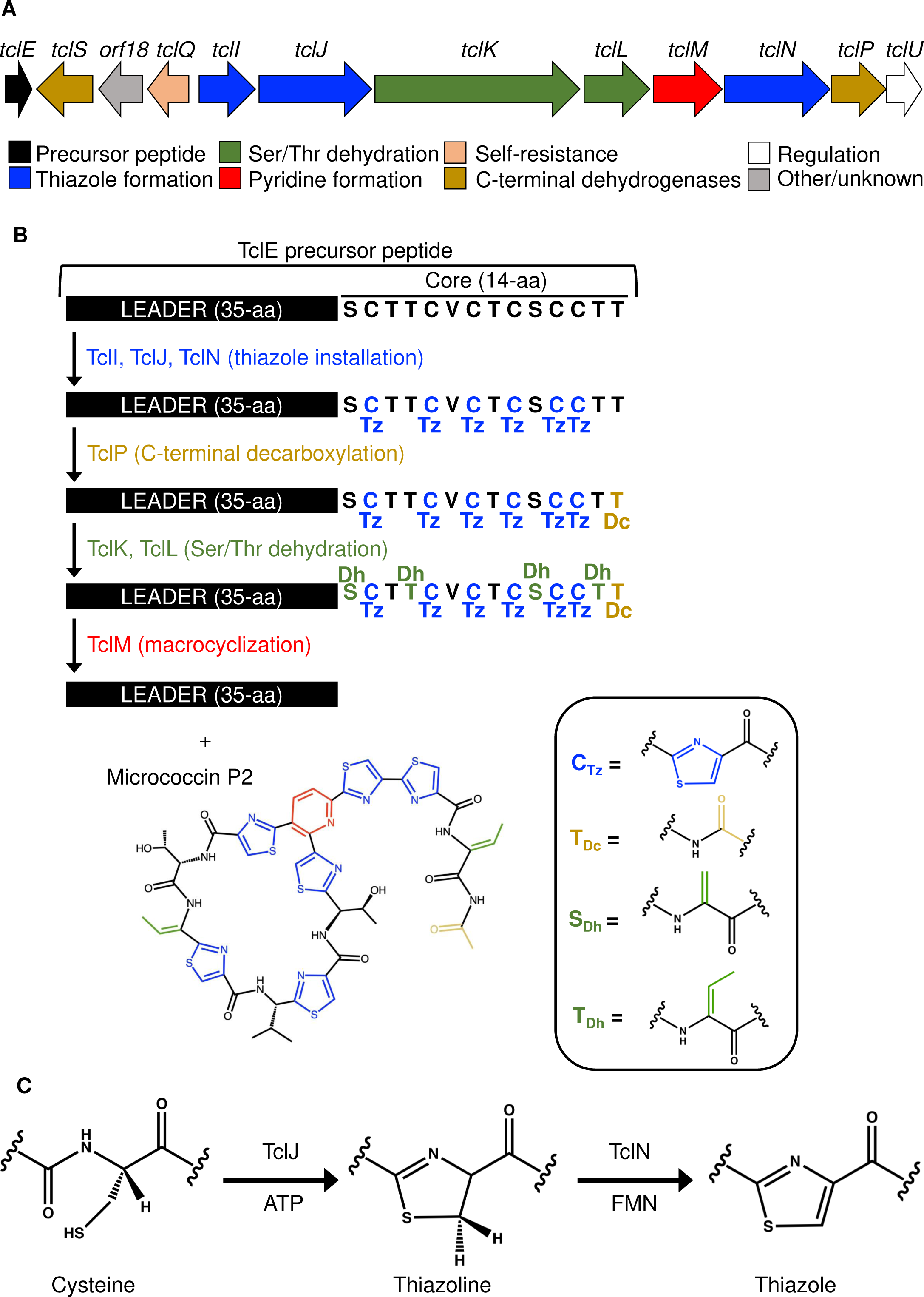
Genes and proteins controlling micrococcin biosynthesis. A. Map of the native *tcl* gene cluster from *M. caseolyticus*. Essential proteins for complete micrococcin production are annotated by colored blocks at the bottom. The gene encoding the precursor peptide (TclE) is colored black, and the genes encoding proteins for thiazole installation (TclI, TclJ, TclN) are colored blue. B. Overview of the micrococcin biosynthetic pathway that converts the TclE core peptide into micrococcin. Modifications and corresponding enzymes are color coded. Abbreviations: Tz, thiazolyl; Dc, decarboxyl; Dh, dehydro. C. Two-step conversion of TclE Cys residues to thiazole by the enzymes TclJ, and TclN.

In this study, we investigate how the thiazole installation proteins in the micrococcin biosynthetic pathway interact with the substrate peptide, and with each other, to carry out these modifications. By conducting a truncation analysis on the TclE leader peptide sequence, we determined a 20-amino acid minimal recognition region required for thiazole installation. Furthermore, by using computational modeling and an *E. coli*-based expression system for mutagenesis and copurification experiments, we demonstrate complex formation between TclI, TclJ and TclN, and we propose a mechanism for cysteine to thiazole conversion in which the scaffold protein TclI coordinates thiazole installation by presenting the TclE substrate to each modifying enzyme in dynamic equilibrium.

## RESULTS

### TclE, TclI, TclJ, and TclN can be functionally expressed in *E. coli*

We engineered a system in which *E. coli* would express codon-optimized *tcl* genes encoding TclE, TclI, TclJ, and TclN. Each of these was engineered with affinity tags in a manner which was previously shown to preserve functionality (31). To test the functionality of *E. coli*-expressed Tcl proteins, we evaluated the *in vitro* conversion of the six TclE Cys residues to heterocycles by mass spectrometry (**Fig. 3**). TclE was purified with an N-terminal cleavable GST tag, and the three other Tcl proteins were purified as complexes using N-terminally His-tagged TclI (these complexes are shown in figures below). In the absence of modifying enzymes, TclE purification and proteolytic removal from the GST tag yields a leader-plus-core fragment of the expected molecular weight (m/z = 5373). When treated with *E. coli*-produced TclI and TclJ (TclN excluded), TclE resolved to two major peaks of m/z=5285 and 5266, consistent with the appearance of 5 and 6 thiazolines, respectively. When treated with *E. coli*-produced TclI, TclJ and TclN, the prominent TclE product has m/z = 5253, consistent with the complete 6-thiazole product with an expected -120 Da change (-6 × (H_2_O+2H)) compared to the unmodified peptide.

**Figure 3.**
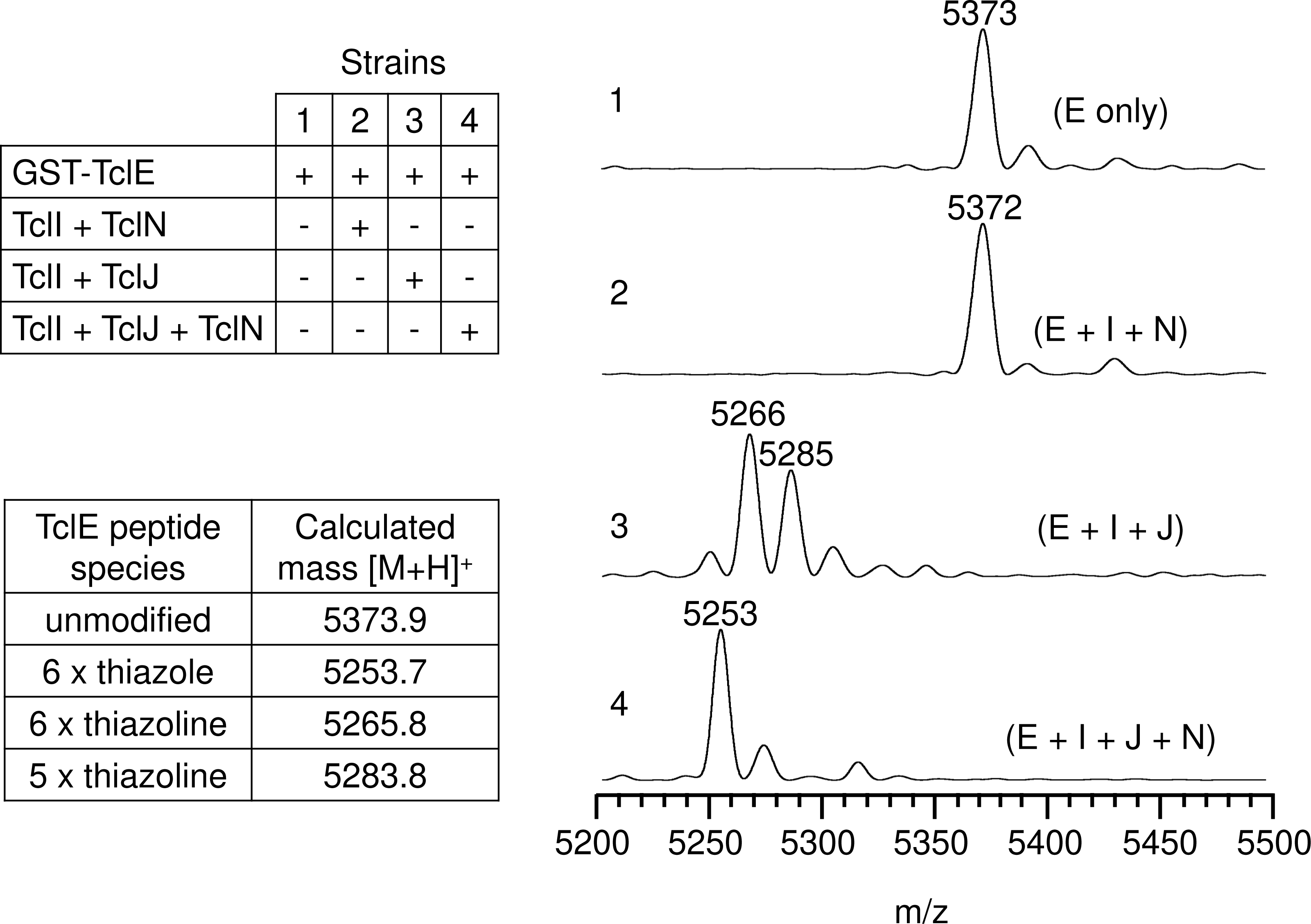
Characterization of functional expression of Tcl TOMM proteins from *E. coli*. MS analysis (MALDI-TOF) of thiazole installation on TclE using purified components from *E. coli*. Reactions included TEV protease to cleave the leader-core region of TclE from the GST tag prior to mass spectrometry analysis. Numbers 1-4 on the left side of each MALDI-TOF spectra represent each of the four strains we used to express the Tcl proteins.

### TclI_NTD_ directly interacts with TclE

We hypothesized that TclI plays a role in TclE recognition. Bioinformatic analysis using the program HHpred (36–38) indicates that TclI is structurally similar to Ocin-ThiF proteins. We obtained a structural model of the TclI:TclE dimer using AlphaFold2 (**Fig. 4A**). The TclI model contains two distinct domains, the N-terminal domain (NTD) is a wHTH structure comprised of three α helices and three β strands, consistent with an RRE. This model places TclE at the interface between the α helices and the 3-stranded β sheet with the leader sequence acting as a fourth β strand, similar to what has been shown for crystallography solved structures in other TOMM systems (20, 22, 24). Key TclE residues in this interaction start with F17 occupying a hydrophobic pocket at the interface of TclI helix 3 and β3. Other predicted key interactions between the TclE leader and the TclI RRE involve three salt bridges: TclE(E21)-TclI(K31), TclE(E22)-TclI(K78), and TclE(E28)-TclI(K22) (**Fig. 4A**). To further investigate whether the TclI_NTD_ is an RRE, we tested for TclI_NTD_ binding to TclE by copurification. We co-expressed His-tagged TclI_NTD_ (residues 1-85) with GST-tagged TclE in *E. coli* and carried out copurification with nickel-NTA beads. As shown in **Figure 4B**, TclI_NTD_ pulls down TclE (**Fig. 4B, Lane 3**), indicating a non-covalent interaction between the two proteins. Furthermore, TclI_NTD_ copurifies the TclE leader region when the TclE core region is absent (**Fig. S1**). We then attempted to determine a minimal TclE leader that interacts with TclI_NTD_, so we generated a series of leader truncations in which 3 residues were consecutively removed from the N-terminus of TclE (**Fig. 5A**) for a total of six TclE truncation variants: Δ3, Δ6, Δ9, Δ12, Δ15, and Δ18. Each truncated variant was co-expressed with His-tagged TclI_NTD_, and nickel pull-down experiments were carried out. As shown in **Figure 5B**, some copurification could be detected for all TclE truncations, but the interaction appears weakened beyond the Δ9 truncation. We observe that, as more of TclE is removed, the amount of TclI_NTD_ purified becomes less. This suggests that the stability of this putative RRE domain is enhanced by the presence of a fully functional leader peptide.

**Figure 4.**
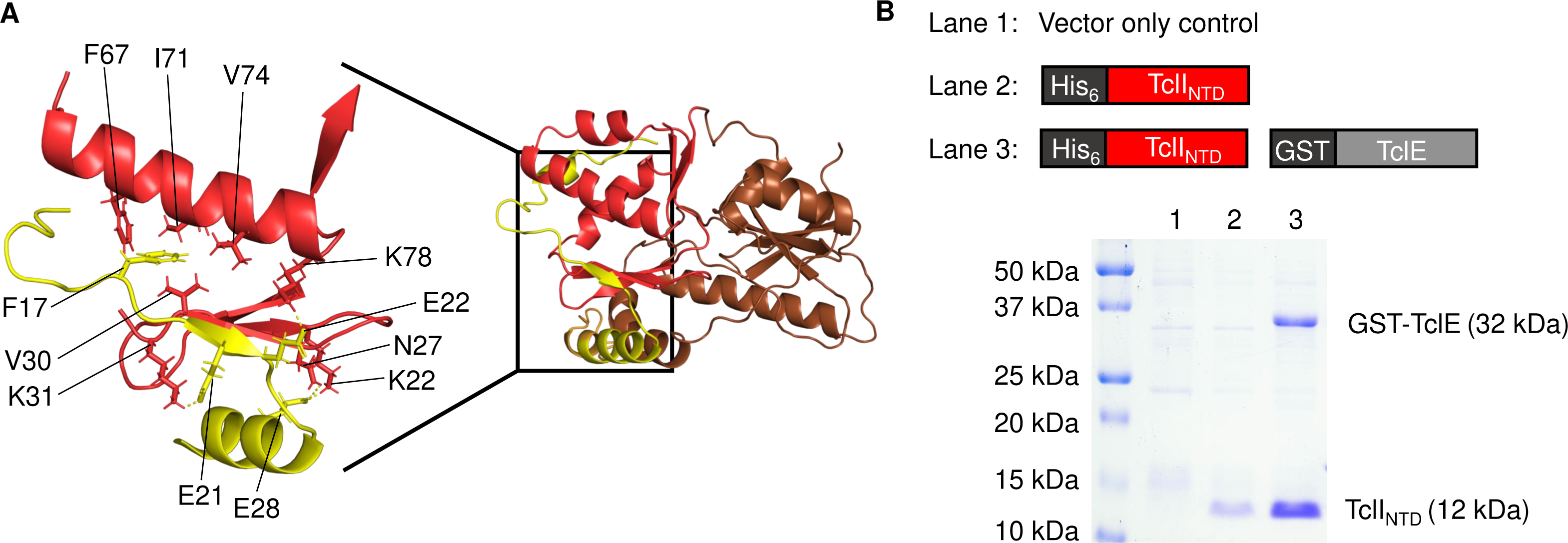
RRE of TclI interacts with TclE. A. AlphaFold2 model of the TclE:TclI dimer. TclE is shown in yellow. TclI_NTD_ is highlighted in red, while TclI_CTD_ is shown in brown. Residues featuring key protein interactions are labeled as follows: TclI_NTD_ (residues F7, V30, V31, I74, V74, K78, N27, K22) in red, and TclE (residues F17, E21, E22, E28) in yellow. B. SDS-PAGE analysis with coomassie blue staining to detect TclE:TclI_NTD_ interactions. Protein identities are given on the right side of the gel, and information of samples loaded in each lane is given above the gel.

**Figure 5.**
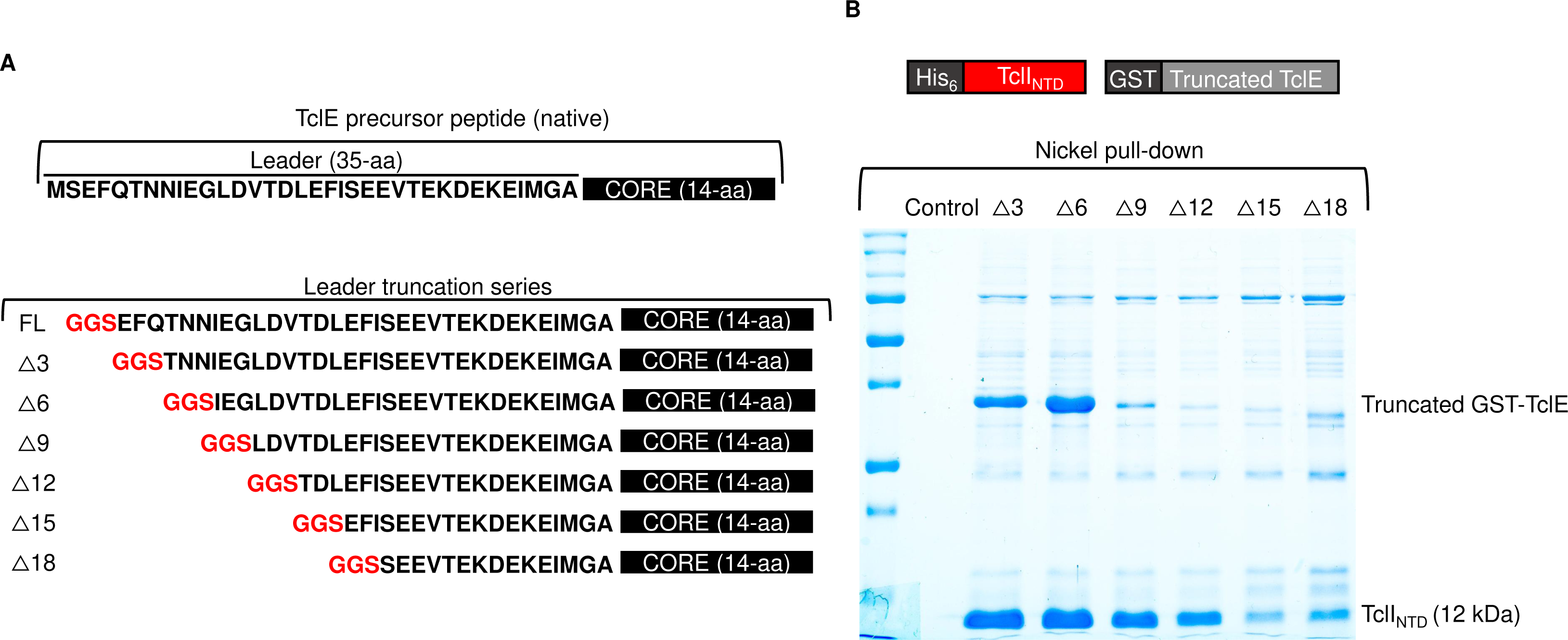
TclE leader truncation analysis. A. Schematic diagram of TclE leader truncations. Native TclE from *M. caseolyticus* is shown on top followed by the mutagenesis analysis on the N-terminus of the TclE leader. B. Nickel copurification experiment to detect TclI_NTD_ interactions with truncated variants of the TclE leader. The top panel shows the Tcl proteins that were co-expressed for this experiment. His-tagged TclI_NTD_ was co-expressed with each TclE leader truncated variant in *E. coli* and subjected to purification with nickel-NTA beads and SDS-PAGE. Protein identities are given on the right side of the gel. For each GST-TclE truncation variant, the molecular weights are as follows: Δ3 (31.4 kDa), Δ6 (31.0 kDa), Δ9 (30.7 kDa), Δ12 (30.4 kDa), Δ15 (30.1 kDa), Δ18 (29.7 kDa).

### Determination of a functionally minimal TclE leader peptide

We then used the TclE truncation series to investigate leader sequence requirements for thiazole installation in *E. coli* cells co-expressing TclI, TclJ, and TclN. We used four of the TclE truncated variants (Δ9, Δ12, Δ15, and Δ18) and assessed thiazole installation by Orbitrap liquid chromatography mass spectrometry (LC-MS) after GST-TclE purification from co-expressing cells (**Fig. 6**). For this analysis, we define “fully modified peptides” as those in which all Cys residues are converted to thiazoles. For the Δ9 truncation 100% of detected TclE peptides were fully modified. The Δ12 variant also yielded products consistent with having a fully modified core, while the Δ15 variant produced a slightly reduced yield of fully modified product (mean= 99.8 ± 0.2%; n=3), with some detected peptides containing intermediates with 4-5 thiazoles. The Δ18 leader truncation yielded multiple intermediates containing a combination of thiazoles, thiazolines and cysteines. The mean of fully modified TclE with the Δ18 truncation is 71.6 ± 4.8% (n=3). We conclude from this that the first 12 N-terminal amino acids of TclE are not required for thiazole installation, and amino acids 12-15 have minimal impact. When amino acids 15-18 are removed, it significantly impairs thiazole installation, potentially due to loss of key TclE-RRE non-covalent interactions (see Fig. 4A). Recall that TclE(F17) was already predicted to mediate an important interaction in a hydrophobic pocket of the modeled TclI_NTD_. Previous studies with TOMM systems such as streptolysin and microcin B17 have shown that the leader peptide is primarily engaged through a conserved FXXXB (B= V, I, or L) motif (39, 40). The TclE leader contains a similar motif (FXXXXB) in residues 17-22. The convergence of these empirical findings with the AlphaFold2 structural prediction and these observations in other TOMM systems gives credibility to a model in which the TclI_NTD_ functions as a genuine RRE in micrococcin biosynthesis.

**Figure 6.**
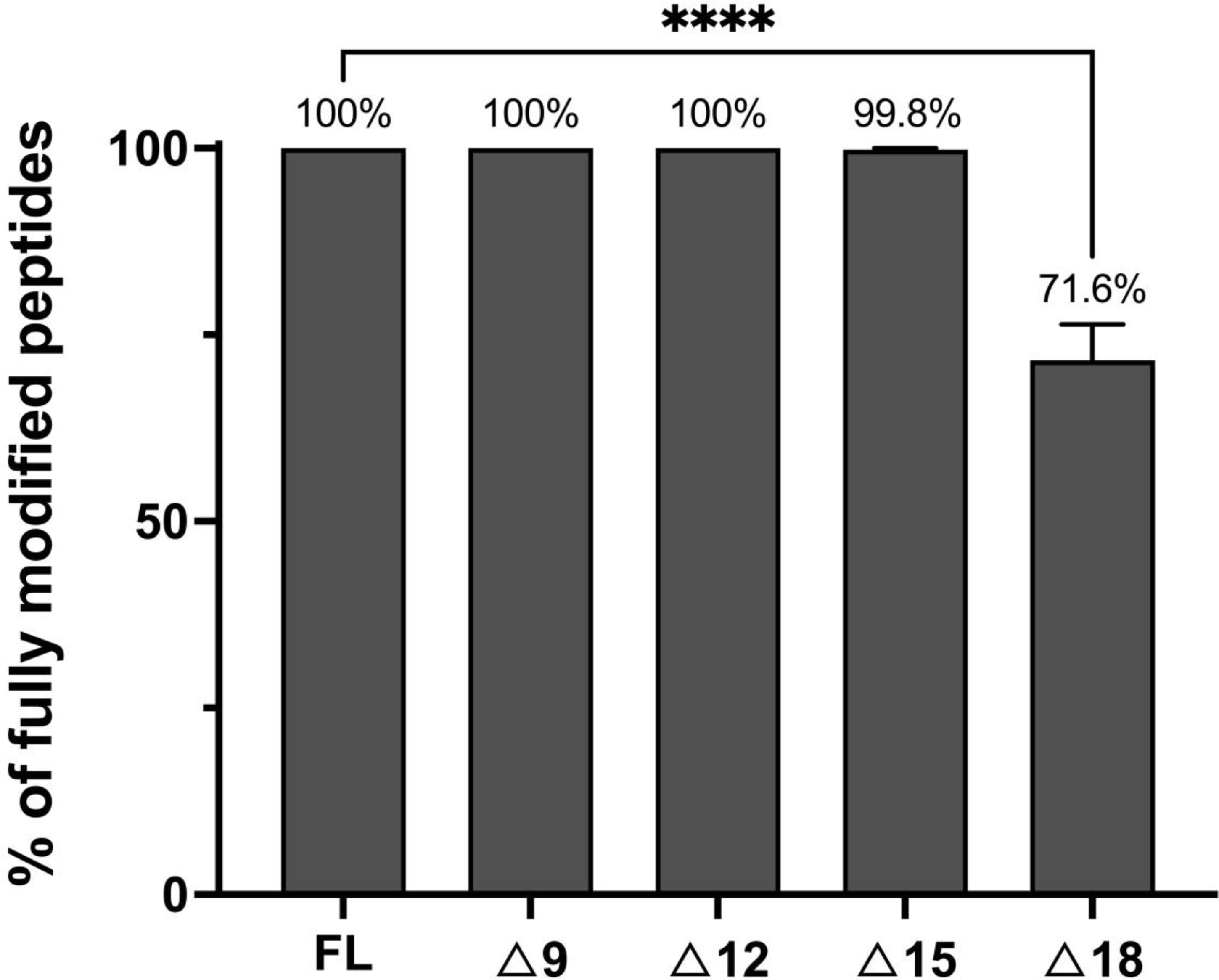
Effects of TclE leader truncations on the production of fully modified core peptides. Fully modified TclE is defined by the presence of six thiazoles in the core peptide. Each bar shows percentage of TclE peptides detected by LC-MS that are fully modified. Each TclE truncated variant (see Fig. 4A; Δ9, Δ12, Δ15, and Δ18) was co-expressed with TclI, TclJ, and TclN in *E. coli* and purified for LC-MS. Reactions included TEV protease to cleave TclE from the GST tag prior to LC-MS analysis. Data are shown as the mean of three independent replicates with standard deviation. Asterisks show statically significant differences (P < 0.05) according to a parametric t-test carried out with the Benjamini, Krieger, and Yekutieli method (57).

### TclI_CTD_ binds to TclJ and TclN

Given that TclI_NTD_ engages with TclE, we hypothesized that the C-terminal domain (CTD) of TclI may be primarily involved in recruiting the enzymatic proteins TclJ and TclN. Ocin-ThiF proteins like TclI have previously been shown to mediate interactions with TOMM enzymes (3, 23, 26). To test interactions of full-length TclI to the modifying enzymes, we co-expressed His-tagged TclI with TclJ or with TclN in *E. coli* and carried out nickel copurification experiments. As shown in **Figure 7** (Lane 3), TclI interacts with both TclJ and TclN when all three proteins are expressed together. When co-expressed with each individual enzyme, TclI also copurifies them (**Fig. 7, Lanes 4-5**). His-tagged TclI_CTD_ (residues 85-242) also copurifies TclJ and TclN (**Fig. 7, lane 7-8**), though TclI_CTD_:TclN shows a weaker interaction than TclI_CTD_:TclJ. These results point to TclI_CTD_ as being sufficient for binding to both TclJ and TclN.

**Figure 7.**
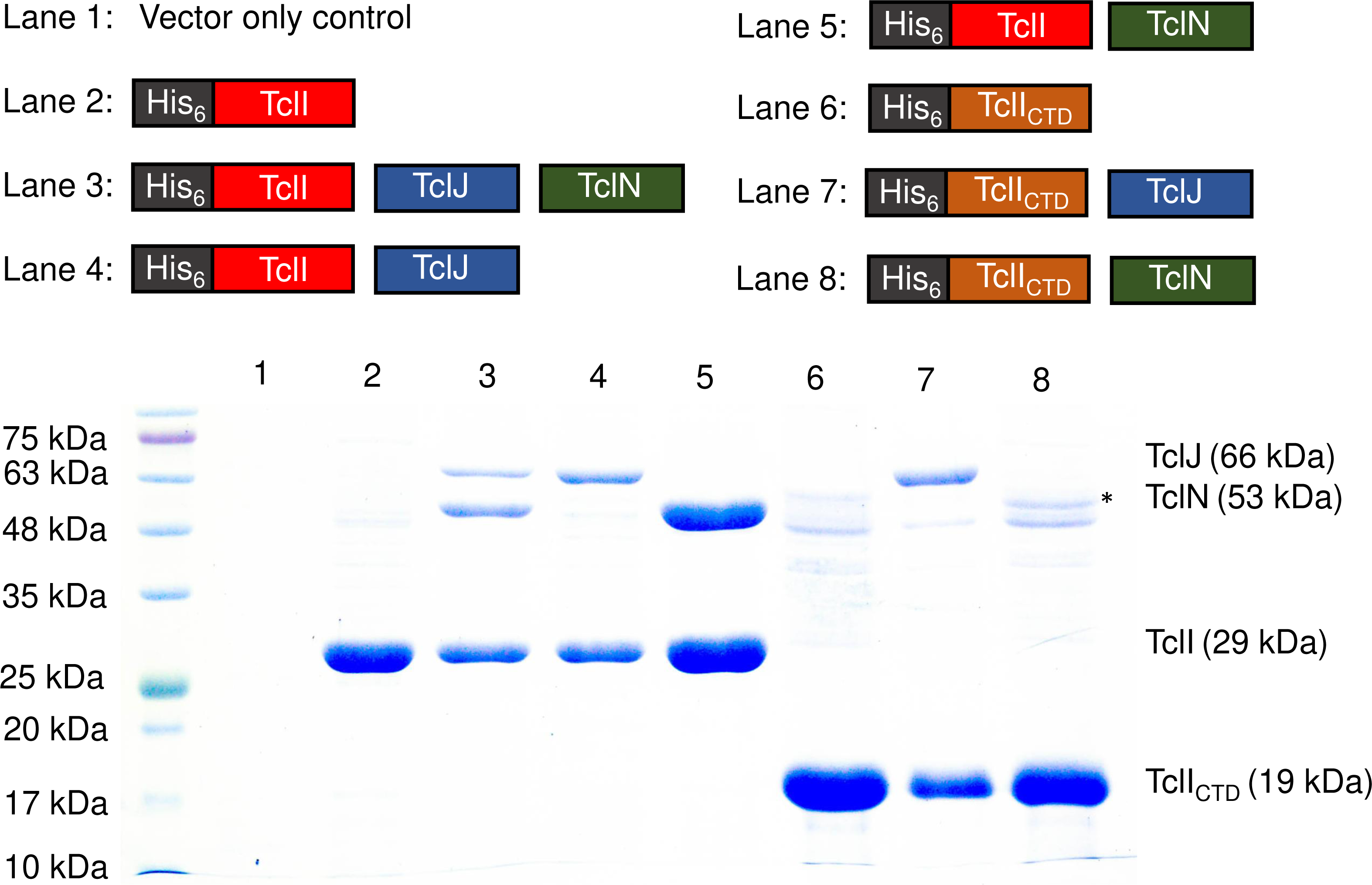
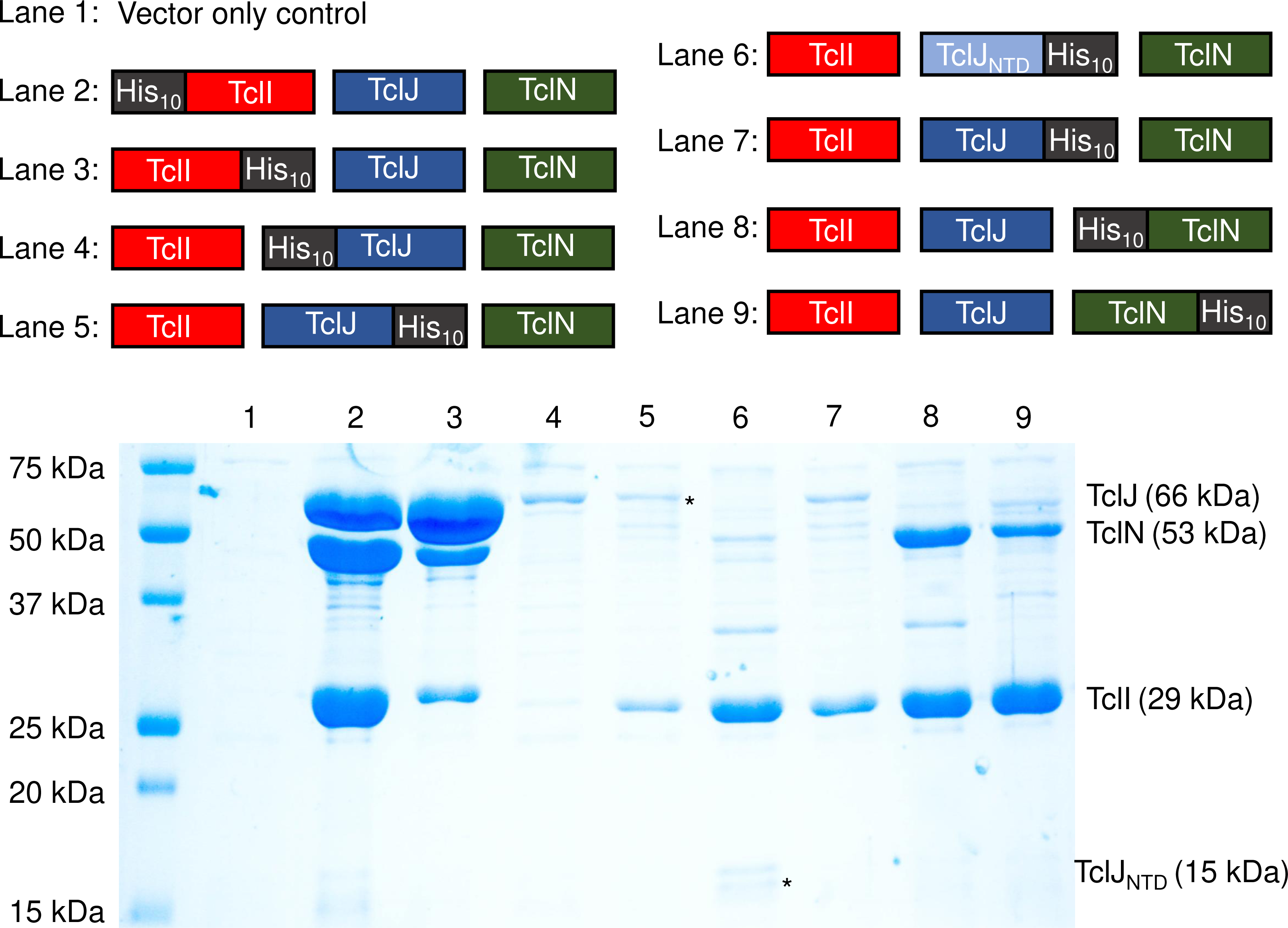
Domain analysis of TclI binding to TclJ and TclN. SDS-PAGE analysis to detect Tcl protein expression and copurification. Maps of His-tagged TclI, TclI_CTD,_ TclJ and TclN used in this study are shown above the gel. Asterisk shows a weak band corresponding to TclN that does not appear in the vector only control (Lane 1) but consistently appears in independently replicated gels. The weak nature of this band could be explained by TclN being degraded by proteases or weak interactions with TclI_NTD_.

Initially, we interpreted these results to mean that TclI has two unique interaction surfaces on its CTD, simultaneously recruiting TclJ and TclN as a stable enzymatic complex. To test whether the TclIJN proteins form a ternary stable complex, we carried out copurification experiments with varying tagging arrangements. We created nine strains expressing different combinations of His-tagged TclI, TclJ and TclN. These experiments show that TclI copurifies TclJ and TclN regardless of whether TclI is N-terminally or C-terminally His-tagged (**Fig. 8, Lanes 2-3**), although TclI may be more poorly expressed or less stable when C-terminally tagged. Furthermore, TclI binding to TclJ appears to be favored when TclI is C-terminally tagged. N-terminally tagged TclJ does not copurify any detectable amounts of TclI or TclN (**Fig. 8, Lane 4**); however, when the tag is moved to the TclJ C-terminus, TclJ robustly pulls down TclI, with little evidence of TclN copurifying (**Fig. 8, Lane 5**). When an N-terminal region of TclJ (residues 1-115) is C-terminally tagged, it very robustly copurifies TclI, and a minor band consistent with TclN can be observed (**Fig. 8, Lane 6**). TclN strongly copurifies TclI regardless of the location of the tag (**Fig. 8, Lanes 8-9**), with possibly a minor TclJ copurification product. In all cases where TclI is pulled down by a tagged version of TclJ or TclN, the band intensities indicate a stoichiometric excess of TclI ranging from 2 to 8 based on densitometry that accounts for staining intensity and molecular weight. In all cases where TclI is pulled down by either tagged enzyme, copurification of the untagged other enzyme is absent or barely evident. From these overall results, we hypothesize that TclI engages with TclJ and TclN in a competitive fashion, and that there may be a weak interaction between the two enzymes.

**Figure 8.**
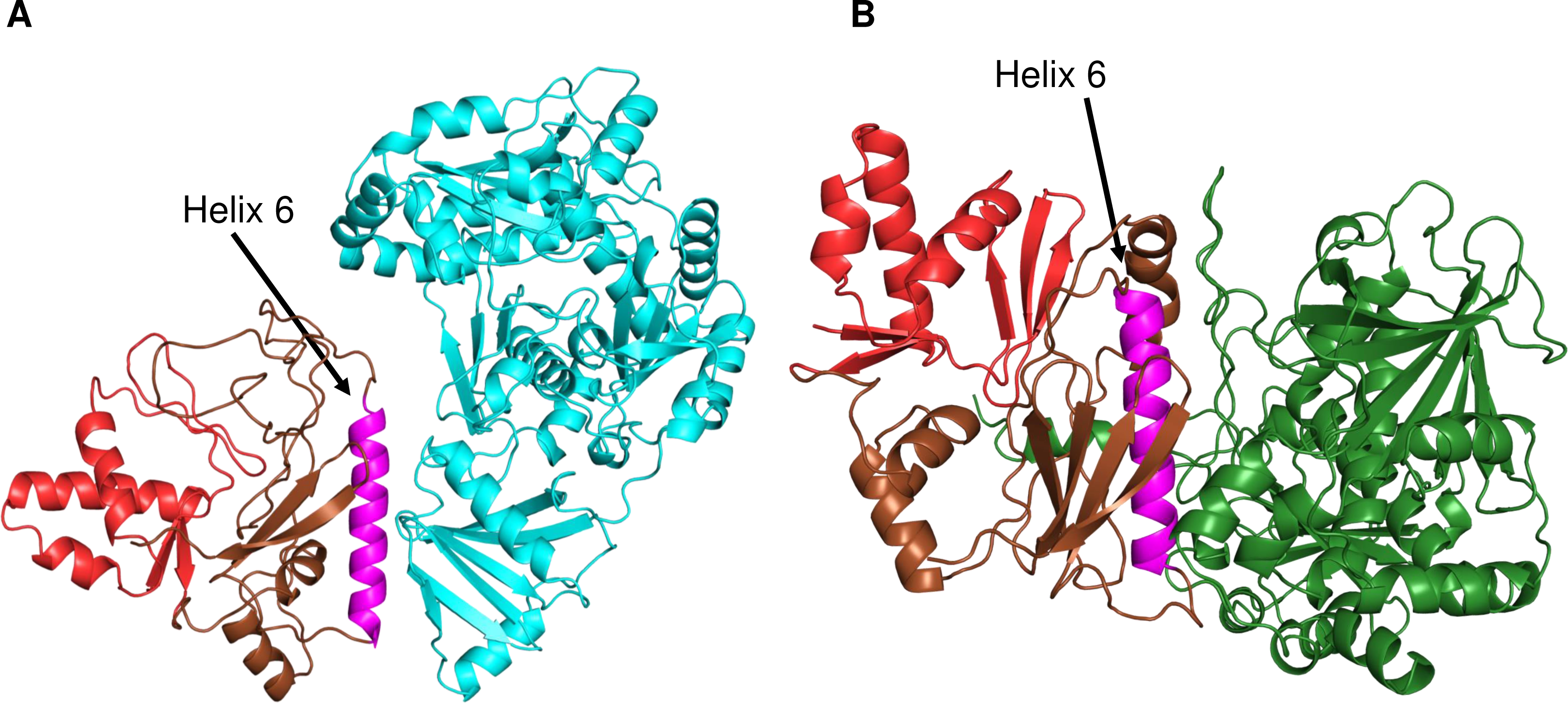
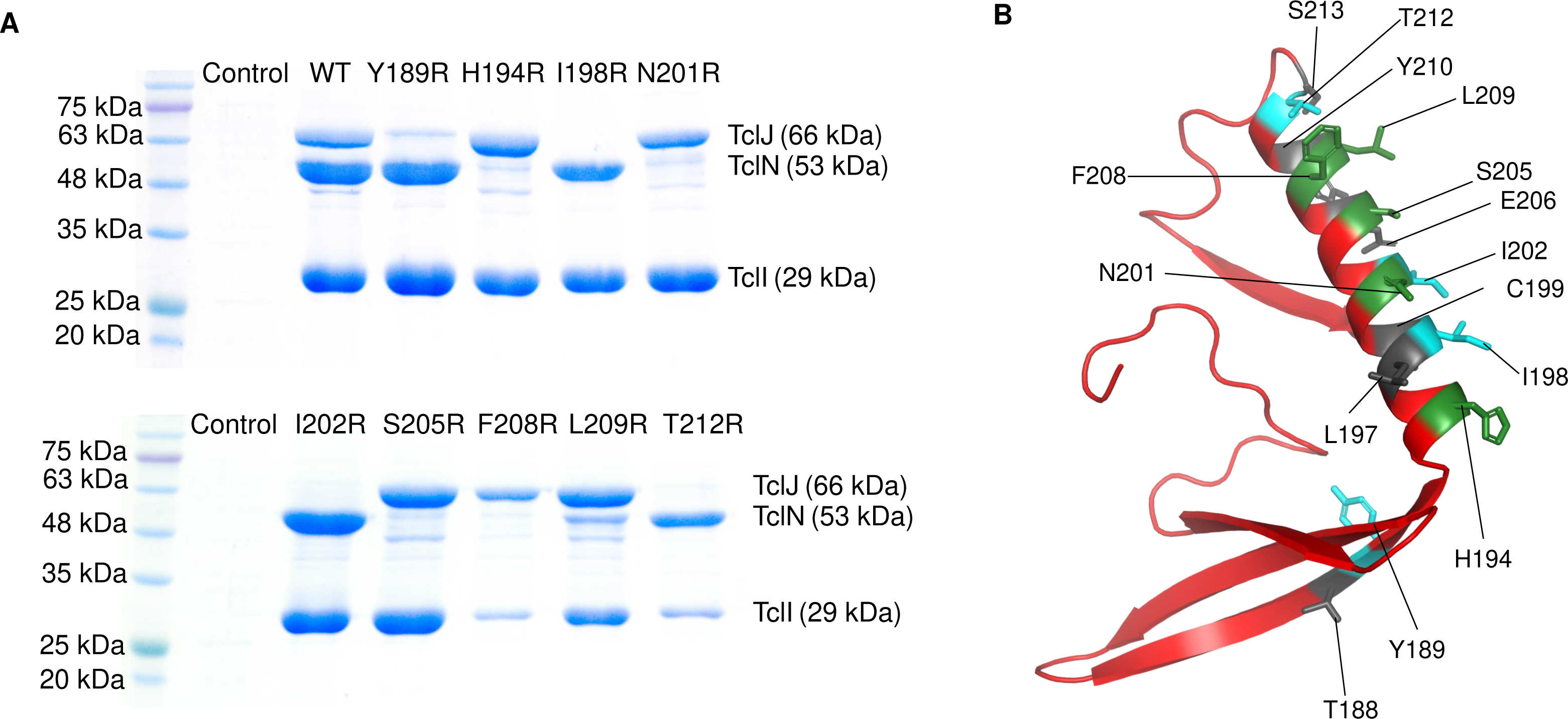
Analysis of TclI, TclJ and TclN interactions as a stable complex. Coomassie-stained SDS-PAGE of purified TclIJN proteins expressed together in *E. coli* to detect complex formation. The upper panel shows the combinations of each his-tagged Tcl protein. The difference between Lane 5 and Lane 7 is that, in Lane 7, the linker between TclJ and the histidine tag is longer (6-Gly) compared to the normal linker (Gly-Gly-Ser) used in Lane 5 and in the other Lanes. Asterisks denote weak protein bands that do not appear in the vector only control (Lane 1) but consistently show up in replicated copurification experiments.

### One surface of TclI facilitates binding to TclJ and TclN

We generated structural models of TclI:TclJ and TclI:TclN dimers using AlphaFold2 (**Fig. 9A, 9B**). According to these two models, TclI_NTD_ folds in a similar conformation for both, while TclI_CTD_ takes on a slightly different structure in each predicted complex, with Helix 6 (TclI residues 188-213) being a central structure for binding to both enzymes. These models reinforce the notion that the TclI:TclJ and TclI:TclN complexes are alternative and mutually exclusive structures. To investigate whether Helix 6 is the primary interaction surface of TclI for binding to the enzymes, we genetically dissected the region corresponding to Helix 6 by substituting each of its 15 surface-exposed residues with an arginine residue and tested for binding to TclJ and TclN under the same conditions used for the copurification shown in **Fig. 8** Lane 2. For this, 15 *E. coli* strains were constructed that express TclJ, TclN and His-tagged TclI with its corresponding Helix 6 substitutions (**Fig. S2**). Copurification experiments show that TclI residues Y189, I198, I202, and T212 are important for the TclI:TclJ interaction since these Arg substitutions abolish binding interactions between the two proteins (**Fig. 10A, Lanes 2, 4, 6, 10**). TclI residues H194, N201, S205, F208, and L209 are critical for interaction with TclN, as when these residues are substituted, TclI:TclN interactions are disrupted (**Fig. 10A, Lanes 3, 5, 7, 8, 9**). Substitutions T188, L197, C199, E206, Y210, and S213 had no effect in TclI interactions with either TclJ or TclN (**see Fig. 10B; Fig. S2**). Substitutions T212R and F208R resulted in a weaker TclI band, suggesting that these changes may also have a negative effect on TclI folding or stability. These findings indicate that Helix 6 is a pivotal structure within TclI responsible for recruiting both TclJ and TclN. Key residues of Helix 6 facilitating binding to each enzyme are interspersed within this postulated surface of TclI_CTD_, further supporting the notion of two enzymes competing for one TclI docking site.

**Figure 9.**
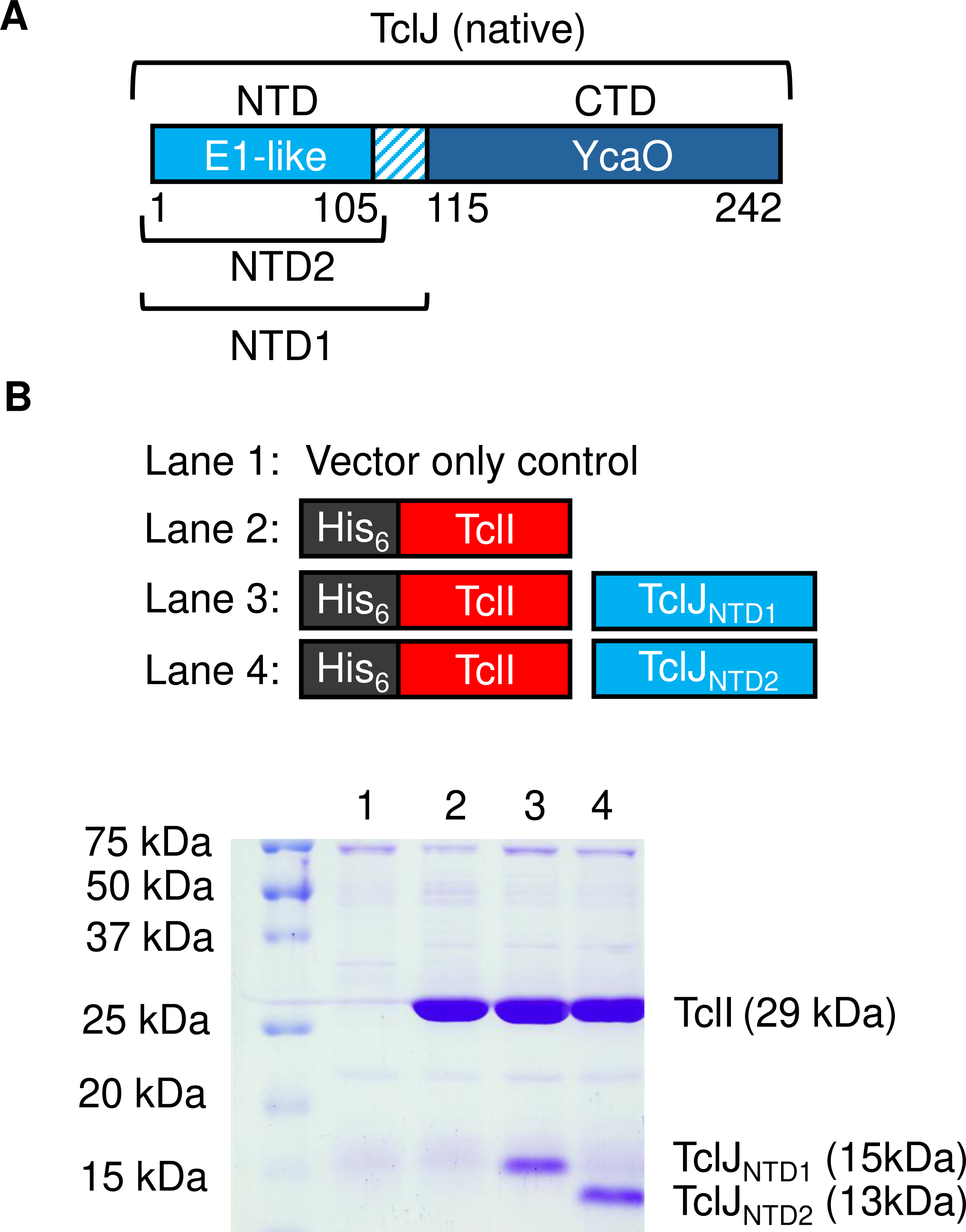
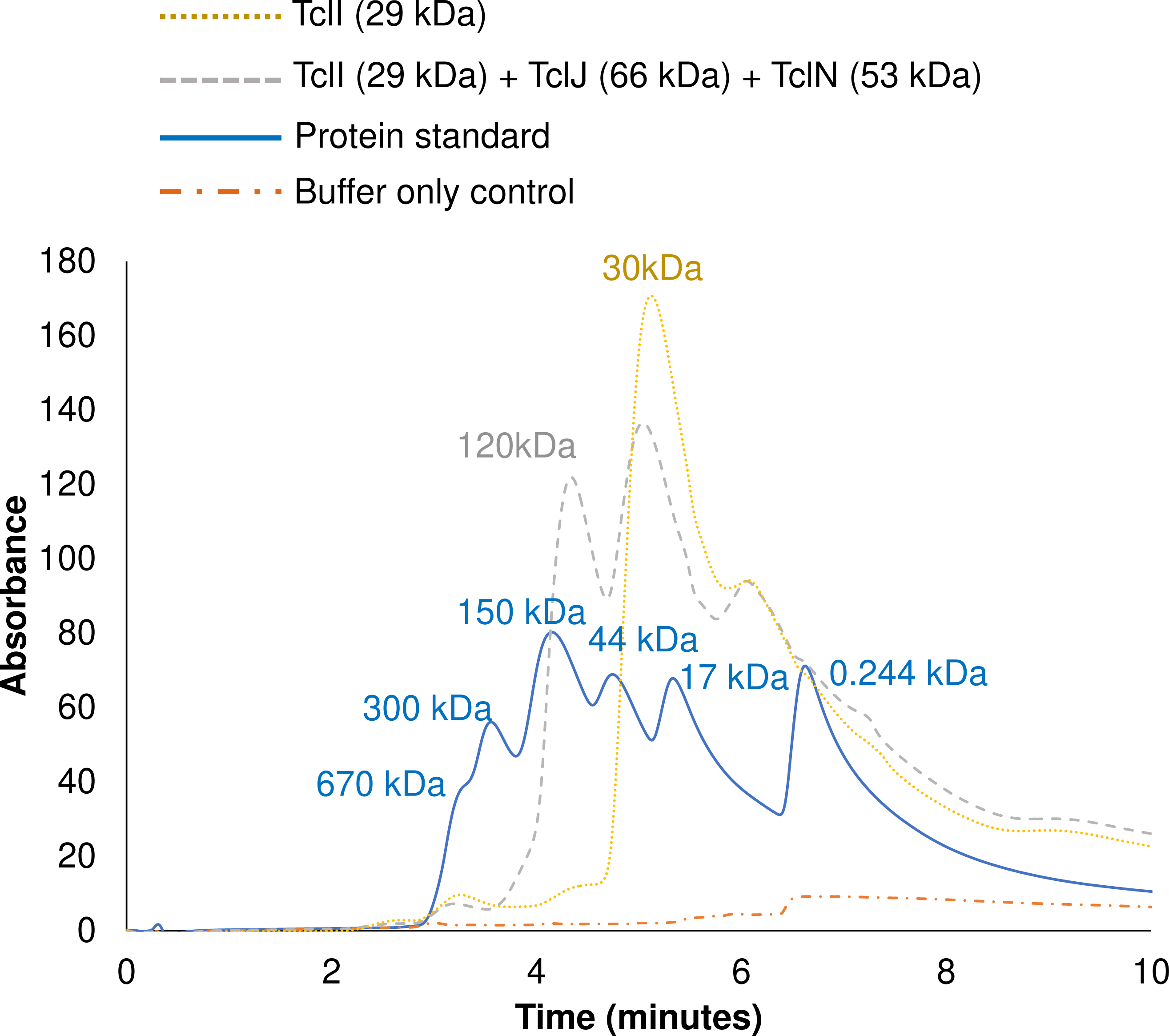
Predicted AlphaFold2 models of the TclI:TclJ (A) and TclI:TclN (B) dimers. TclI_NTD_ is shown in red, and TclI_CTD_ is shown in brown with the central Helix 6 highlighted in magenta. TclJ is depicted in cyan and TclN is shown in green.

**Figure 10.**
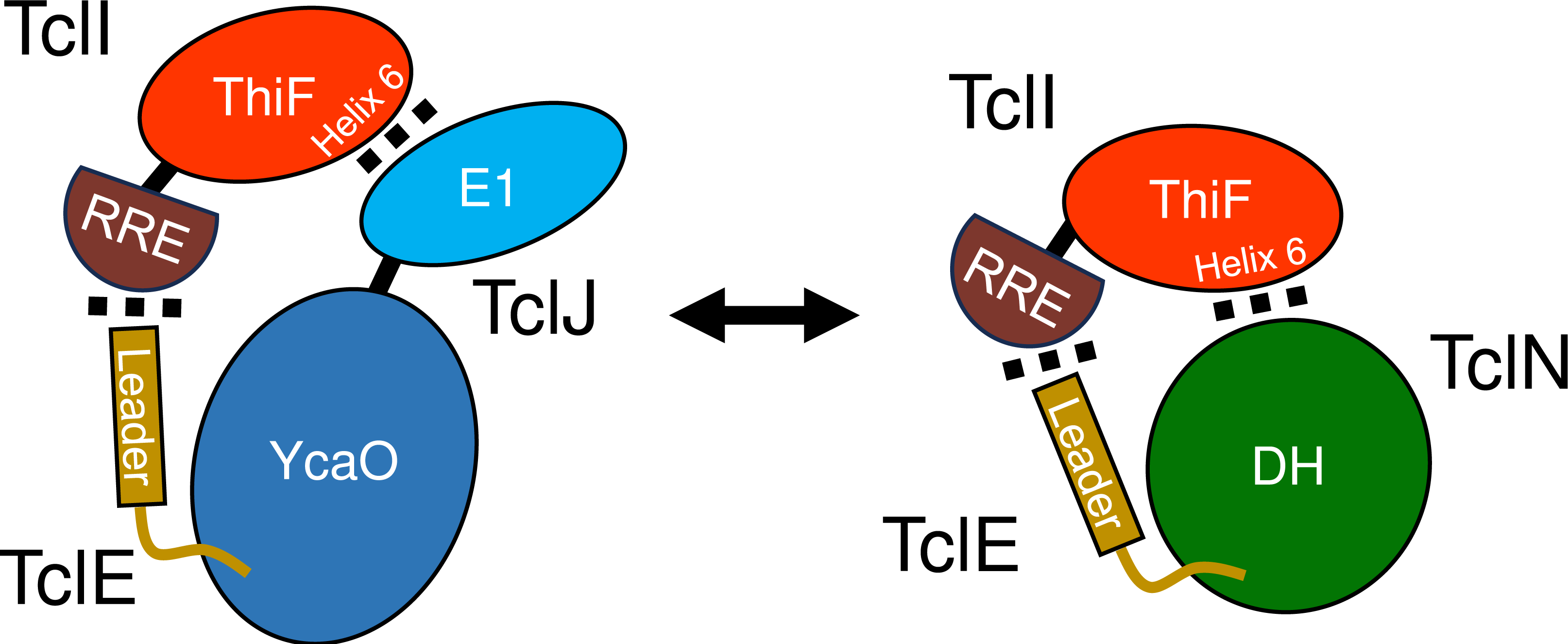
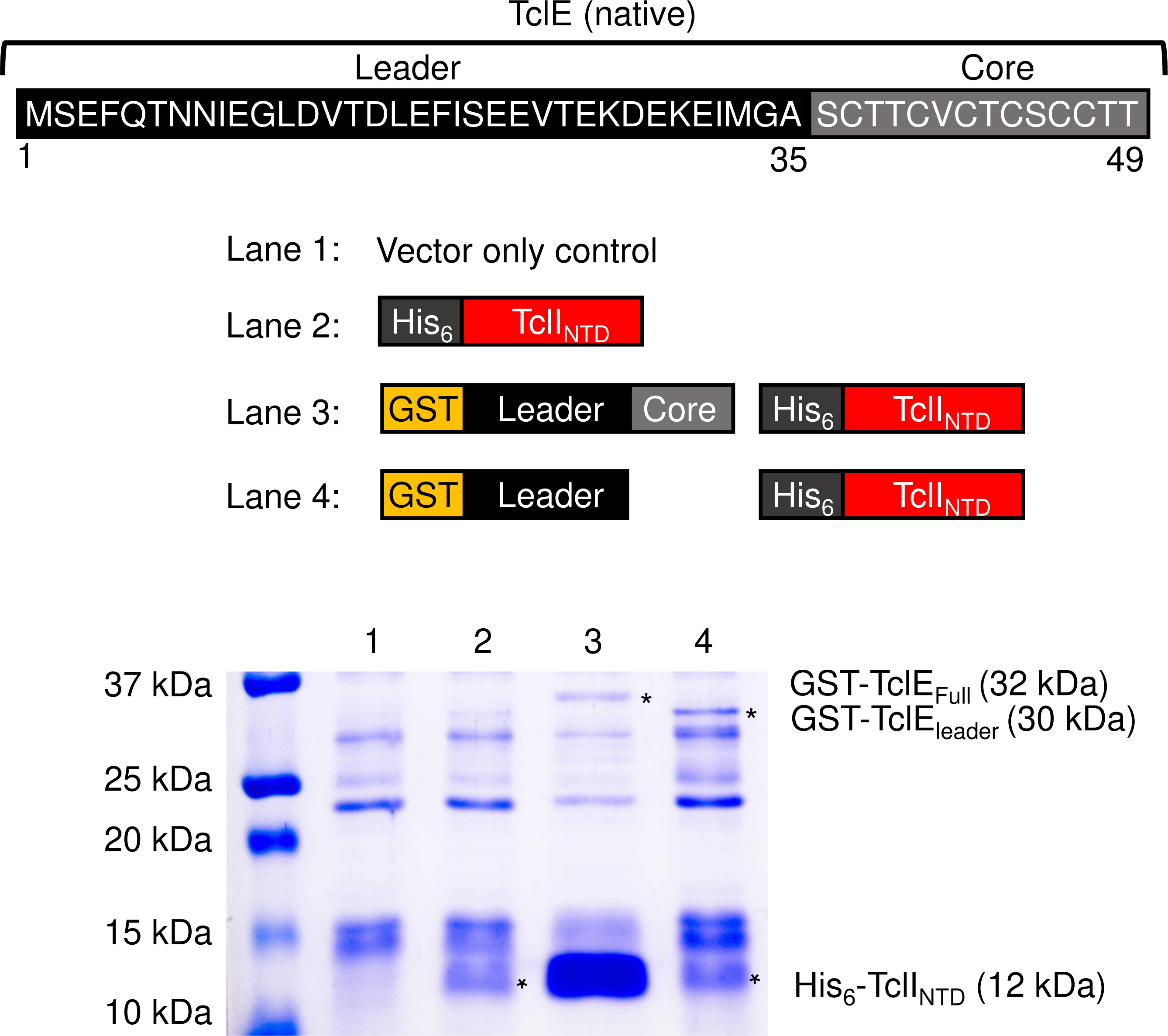
Dissection of Helix 6 on TclI to determine key interactions with TclJ and TclN. (A) Nickel copurification experiment with His-tagged TclI with its corresponding Helix 6 substitution co-expressed with TclJ and TclN. Each substitution on TclI is labeled on each lane. (B) Depiction of Helix 6 highlighting key residues for interacting with TclJ in cyan (Y189, I198, I202, T212) and TclN in green (H194, N201, S205, F208, L209). Residues highlighted in gray (T188, L197, C199, E206, Y210, S213) had no effect in TclI interactions with either TclJ or TclN (see Fig. S2).

### An independent domain of TclJ facilitates binding to TclI

We wanted to further investigate how the N-terminal region of TclJ (when C-terminally tagged), is able to pull down TclI, while the N-terminally tagged full-length TclJ is unable to (see **Fig. 8, Lanes 4-5**). We reasoned that the N-terminus of full-length TclJ is important for TclI interaction. We obtained AlphaFold2 models of TclJ and TclN (**Fig. S3A, and S3B**). The TclJ model features two distinct domains, corresponding to a small NTD (residues 1-105) with the enzyme active site in a larger YcaO-like CTD (residues 115-242). HHpred analysis showed that the NTD of TclJ is a short E1-like domain likely involved in protein-protein interactions. The TclJ C-terminus is embedded in its putative active site, while its N-terminus is predicted to be surface exposed on this E1-like NTD. We hypothesized that TclJ_NTD_ folds independently and facilitates binding to TclI. To test this, two versions of the TclJ_NTD_ (resides 1-105 and residues 1-115) were co-expressed with His-tagged TclI for nickel copurification. Both versions of TclJ_NTD_ copurified with TclI (**Fig. 11, Lanes 3-4**) indicating that the E1-like domain of TclJ (residues 1-105) folds independently and forms a TclI-binding domain. Tags on the N-terminus of this domain appear to obstruct TclI binding (see **Fig. 8, Lane 4**).

**Figure 11.**
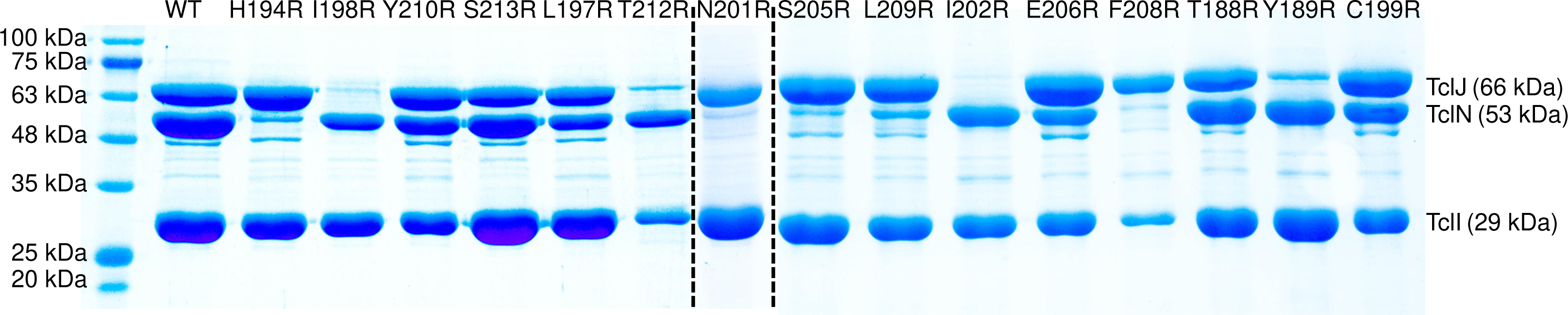
E1-like domain analysis of TclJ. A. Map of TclJ highlighting its E1-like and YcaO domains. Labeled are the two NTD fragments of TclJ used in this study. B. SDS-PAGE analysis to detect interactions between the two TclJ_NTD_ fragments (NTD1, and NTD2) and TclI.

### Analysis of TclIJN complexes by size-exclusion chromatography

To investigate this model, we evaluated the TclIJN complexes using size-exclusion chromatography. We first established the oligomeric state of purified TclI, which was consistent with a monomer (**Fig. 12**). However, when TclI, TclJ and TclN were analyzed as a mixture, a new peak arose with a retention time consistent with a molecular weight of ∼120 kDa, suggesting molecular units that contain two copies of TclI and one copy of either enzyme (2TclI:1TclJ, expected: 121 kDa or 2TclI:1TclN, expected: 111 kDa) (**Fig. 12**).

**Figure 12.**
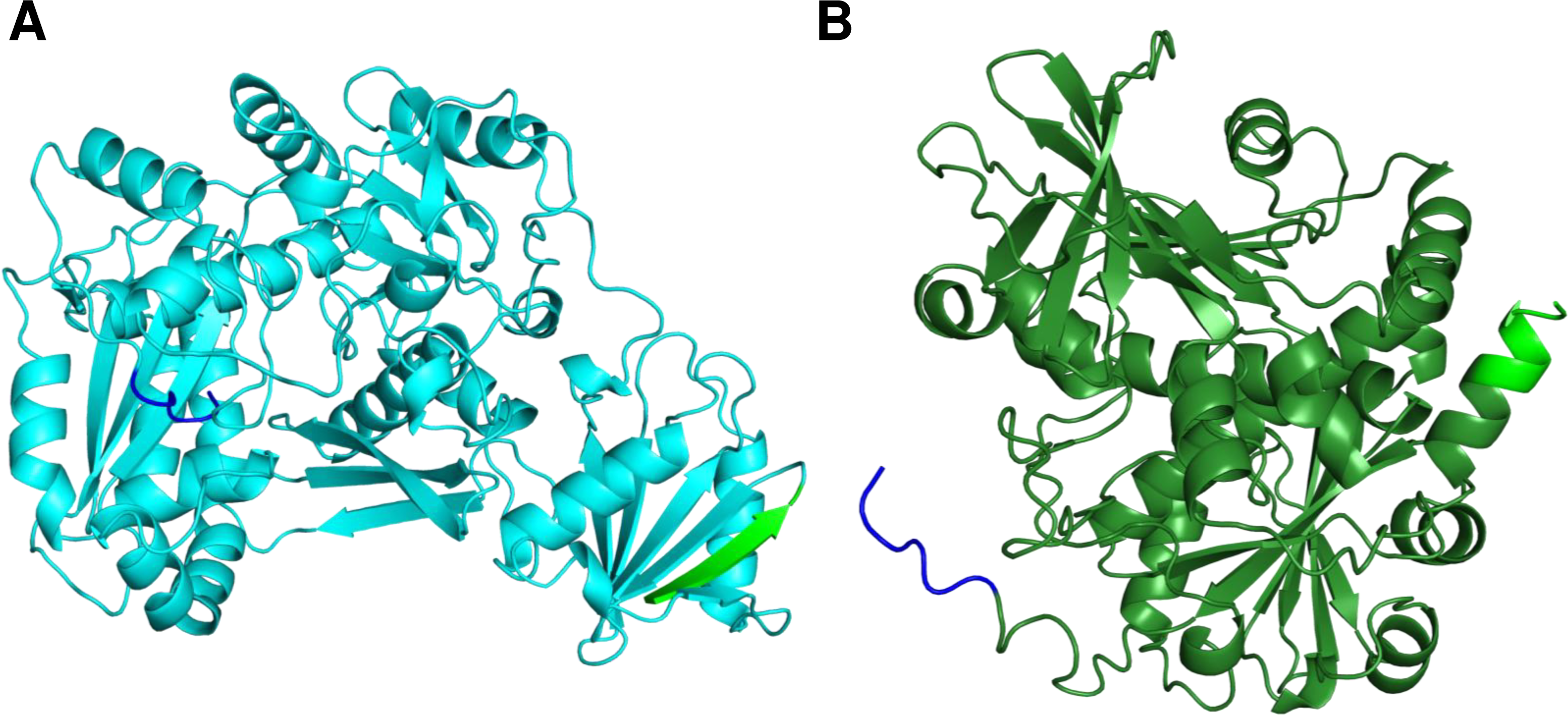
Size-exclusion chromatography of purified TclI and TclIJN. Yellow dotted line indicates chromatogram of purified TclI. Grey dotted line shows the chromatograph for the TclIJN purified complex. The blue chromatogram represents the protein ladder, and the dotted orange line shows the chromatogram for the buffer only control.

## DISCUSSION

The Tcl biosynthetic pathway from *M. caseolyticus* synthesizes micrococcin from a precursor peptide (TclE) by orchestrating the following post-translational modifications: Cys-to-thiazole conversion, C-terminal decarboxylation, Ser/Thr dehydration, and macrocyclization. The work provided in this study focuses on the thiazole installation step to provide a better understanding of the complex protein interactions of thiazole modifying enzymatic proteins. Thiazole installation occurs in a two-step process in which a cyclodehydratase (TclJ) converts six cysteines in the core peptide of TclE into thiazolines, and then a dehydrogenase (TclN) converts the thiazolines into thiazoles. In this study, we determined a minimal region on the leader peptide that is required for thiazole installation, we show complex formation between TclI, TclJ, and TclN, and further define conserved domains within these proteins that play a critical role mediating protein interactions for cysteine to thiazole conversion.

Thiazole/oxazole modified microcins (TOMMs) are derived from a precursor peptide that contains an N-terminal leader and a C-terminal core. Modifications in TOMM precursor peptides are orchestrated by an E1-like scaffold protein, a YcaO cyclodehydratase, and a dehydrogenase; in some cases, a highly divergent Ocin-ThiF-like protein is also present (16, 21, 26, 41, 42). The E1-like and Ocin-ThiF-like proteins are involved with leader peptide binding through an RRE domain that features a winged helix-turn-helix (wHTH) structure comprised of three α-helices and a three-stranded β-sheet (15). The purpose of the leader peptide is to recruit the biosynthetic enzymes, but they have shown variations in their roles in the biosynthesis of diverse TOMMs (20, 43–45). For instance, studies in Lantibiotics such as Nisin have shown that the core can be processed in the absence of the leader peptide, albeit with reduced efficiency (46, 47). Reports for cyanobactin processing by LynD have shown that in the absence of the leader, the core does not get modified; however, if the leader is provided as a separate peptide (*in trans*) or appended to the N-terminus of the biosynthetic enzyme, the substrate is fully modified (20, 43, 48). In this study, we define the NTD of TclI as the RRE that directly interacts with the TclE leader, and we define a minimal leader region that is essential for binding to the RRE and allows for complete thiazole installation by the biosynthetic enzymes. Reports on streptolysin and microcin B17 have determined that the leader peptide is recognized by a conserved FXXXB (B= V, I, or L) motif (39, 40). Our studies show that truncation of a similar motif (FXXXXB) in the TclE leader significantly reduces RRE binding and thiazole installation. This suggest a shared strategy in the leader of TOMM precursors that has been conserved through evolutionary timescale.

TOMM biosynthetic clusters have shown a wide variety of structural variations in how the biosynthetic enzymes interact with each other to install azole heterocycles onto a precursor peptide. The solved structures of the TruD and LynD cyclodehydratases provide an example of fused cyclodehydratases that contain an E1-like and a YcaO domain (49, 50). TOMM proteins for the Heterocycloanthracin (Hca) biosynthetic cluster feature a single copy of an Ocin-ThiF-like protein (HcaF), a fused E1-YcaO cyclodehydratase (HcaD), and a dehydrogenase (HcaB), with the Ocin-ThiF-like protein comprising of an RRE and interacting in a 1:1 ratio with the fused cyclodehydratase; attempts to elucidate interactions between the scafold protein and the dehydrogenase for the Hca pathway have not been possible because the dehydrogenase is heavily proteolyzed when expressed in *E. coli* (26). The Microcin B17 synthetase features a discrete cyclodehydratase (McbD) and dehydrogenase (McbC) that are dependent on the presence of an E1-like protein (McbB) for substrate recognition forming a higher order active complex with a ratio of 4McbB:2McbD:2McbC (22). The TOMM proteins for the biosynthesis of sulfomycin feature combinations of RREs (SulB, SulF), E1/Ocin-ThiF-like domains (SulB, SulC, SulE, SulF), cyclodehydratases (SulC, SulD) and dehydrogenases (SulF, SulG). Combinations of these proteins form three active complexes (SulBC, SulEFG, SulDEFG) that achieve azole formation in the substrate peptide (23). Our findings show that the TOMM proteins from the Tcl biosynthetic cluster comprise of an Ocin-ThiF-like protein (TclI) that includes an RRE, a fused cyclodehydratase (TclJ) that includes an E1-like domain and a YcaO domain, and a dehydrogenase (TclN). TclI binds to the TclE leader through its RRE and presents the core peptide to each enzyme in a mutually exclusive manner, forming two distinct complexes (TclI:TclJ, and TclI:TclN). Our results also suggest that the enzymes may interact weakly with each other creating spatial constraints, so that both enzymes are readily available to bind to TclI when taking turns to modify TclE in a coordinated manner (**Fig. 13**).

**Figure 13.**
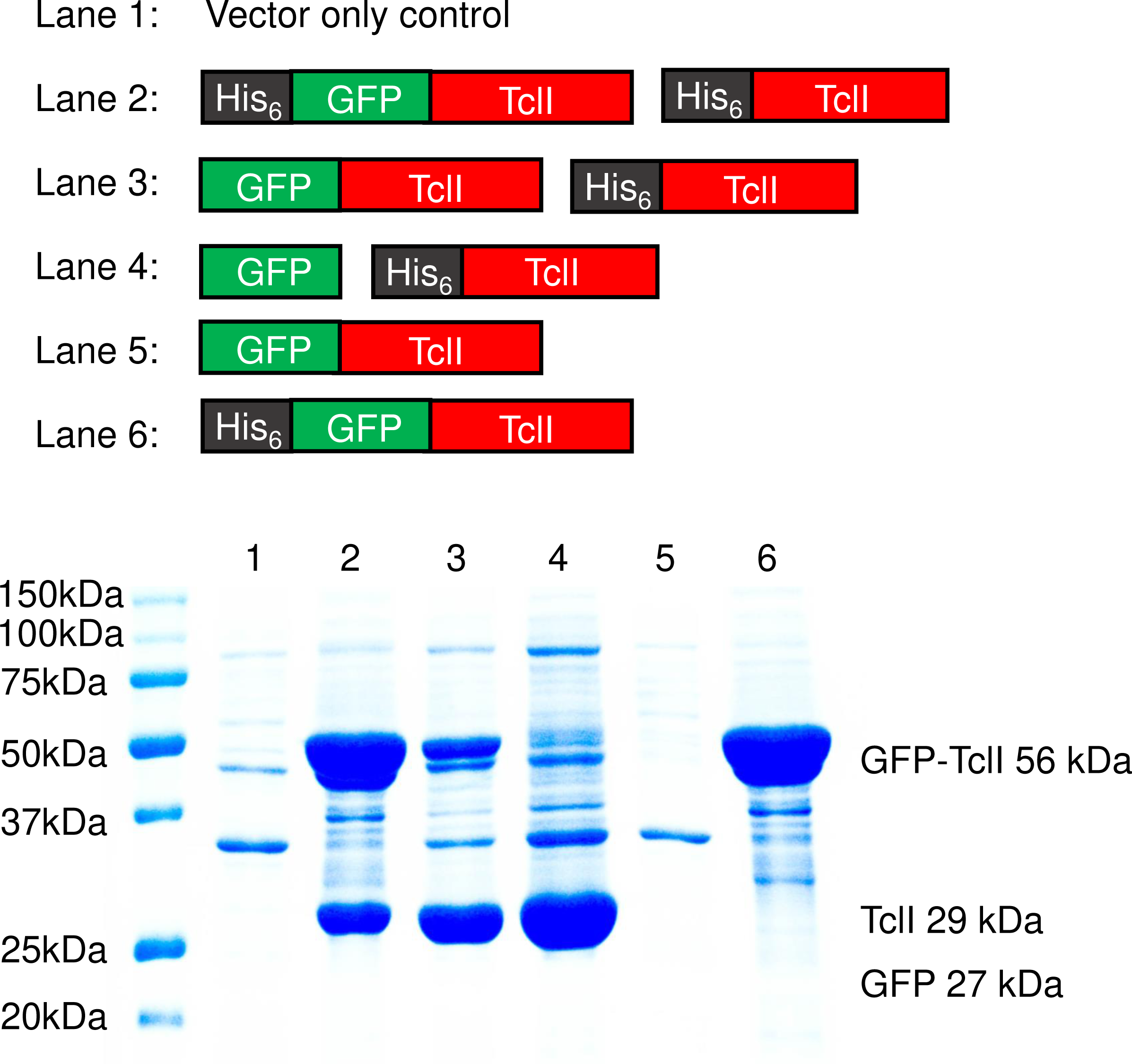
Paradigm for cysteine to thiazole conversion by TclI, TclJ and TclN on the precursor peptide TclE in the biosynthesis of micrococcin. Schematic model of the two functional complexes (TclI:TclJ, and TclI:TclN) that assemble to achieve two-step thiazole installation on the core of TclE.

Size exclusion chromatography data suggests that TclI exists in a monomeric conformation when expressed by itself, but it adopts a homodimeric structure when interacting with the enzymes (**Fig. 12**). Furthermore, preliminary copurification experiments (**Fig. S4**) suggest that TclI may indeed form a homodimer, thus the ratio of the active complexes might consist of 2TclI:1TclJ, and 2TclI:1TclN, but further investigation is required to validate this claim. Overall, the findings from this study have not only elucidated key domains of TOMM proteins in the Tcl biosynthetic cluster, but it provides evidence of the high functional and structural diversity of the TOMM protein family.

## MATERIALS AND METHODS

### Plasmids, strains, and culture conditions

The bacterial strains and plasmids used in this study are summarized in Tables S1 and S2. For full plasmid sequences, refer to the Supplemental Materials file. Plasmids were constructed and maintained in *E. coli* strain DH5α. The *tcl* genes were synthesized with codon optimization for expression in *E. coli*. The sequences of vectors and *tcl* gene inserts are given in the supplemental information file. For protein expression, plasmids were transformed into strain BL21, Nico 21 (DE3) or DH5α. All bacterial cultures were grown in Luria broth (LB: per liter, 10 g Bacto tryptone, 5 g Bacto yeast extract, 5 g NaCl, 1 ml 2N NaOH). Antibiotics used were kanamycin (30 µg/ml), ampicillin (100 µg/ml), and chloramphenicol (30 µg/ml). Cultures were induced for protein expression using 0.3 mM isopropyl β-D-1-thiogalactopyranoside (IPTG).

### Tcl protein expression and purification

To prepare samples for SDS-PAGE analysis, overnight liquid cultures (4 ml) were grown from single colonies in the presence of appropriate antibiotics. 50-ml cultures were inoculated with 2 ml of overnight culture, allowed to grow at 30°C for 1 h, followed by induction with IPTG for another 6 h at 30°C. Cells were collected by centrifuging the culture, and cell pellets were frozen at -80°C for a minimum of 1 h. Cell pellets were then processed for protein purification with either Ni-NTA-linked (for the His_6_ tag) or glutathione-linked (for the GST tag) resin, as detailed below.

For His purification, cell pellets were thawed on ice and re-suspended in 1 ml of lysis buffer (50 mM HEPES pH 7.8, 300 mM NaCl, 0.2 % Triton X-100, 0.5 mg/ml lysozyme, 60 mM imidazole, 1 mM EDTA). Lysis took place for 1 h at 4°C. Cell lysates were then sonicated 4 × 20 sec using a probe sonicator to ensure complete lysis and fragmentation of DNA. Samples were centrifuged at 13,000 rpm for 10 min (4°C) and approximately 1 ml of supernatant was transferred to a new microcentrifuge tube. Supernatant was incubated end-over-end with 50 μl of NTA-nickel agarose beads (Qiagen) at 4°C for 30 min. Nickel beads were pelleted at 13,000 rpm for 30 seconds and washed 3 × 1 ml with wash buffer (60 mM imidazole, 300 mM NaCl, 50 mM HEPES pH 7.8). Purified proteins were then eluted in 50 μl of 2× SDS sample buffer (20% glycerol, 83 mM Tris pH 6.8, 40 mg/ml sodium dodecyl sulfate (SDS), 0.01% bromophenol blue, 0.03 μl/ml 2-mercaptoethanol).

For GST purifications, cell pellets were thawed on ice and re-suspended in 1 ml lysis buffer (50 mM Tris 8.0, 150 mM NaCl, 0.5 mg/ml lysozyme, 2 mM EDTA, and 0.2% Triton X-100). Cells were lysed for 1 h at 4°C then dithiothreitol (DTT) was added to a final concentration of 1.5 mM. Samples were sonicated 4 × 20 sec. Cell lysates were centrifuged at 13,000 rpm at 4°C for 10 min to pellet cell debris. Approximately 1 ml of supernatant was transferred to a new microcentrifuge tube. 50 μl of unwashed glutathione-agarose beads was added and samples were rotated end over end for 45 min at 4°C. Slurry was pelleted at 13,000 rpm for 30 seconds. Supernatant was removed and beads were washed 3 × 1 ml GST buffer (50 mM Tris 8.0, 150 mM NaCl). GST buffer was completely removed, and proteins were eluted from resin in 50 μl of 2× SDS sample buffer.

Purified samples were heated at 100°C for 5 min. Unless stated otherwise, 8 μl of supernatant was loaded onto a 12% resolving Laemmli gel with a 4% stacking gel. Gels were run using 1× Laemmli running buffer, stained overnight in coomassie blue stain, followed by destaining and soaking in water prior to imaging.

### Mass spectrometry analysis of TclE processing

For purification of His_6_-tagged enzymes for mass spectrometry analysis, 25-ml overnight cultures were grown from single colonies. These overnight cultures were then used to inoculate 1 L induction cultures (30°C). After 1 h, IPTG was added and the cultures were grown for an additional 6 hours. The cells were harvested by centrifugation and the cell pellets were frozen at -80°C overnight. For copurification of His_6_-TclIJ, His_6_-TclIN, or His_6_-TclIJN the cells were then thawed on ice with the addition of lysis buffer (50 mM HEPES, 150 mM NaCl, pH 7.8). A protease inhibitor tablet (Roche), 0.2% Triton X-100 and 0.5 mg/ml lysozyme were added and the cells were incubated on ice for 1 h. Complete lysis was achieved by sonication for 2 min on ice using a Branson Sonifier 450, followed by centrifugation for 20 min at 32,539 × g. The supernatant was incubated with 1 ml of Talon resin for 30 min at 4°C. Resin was washed with 3 × 10 ml lysis buffer, followed by elution with lysis buffer plus 75 mM imidazole (4 × 1 ml). The elution fractions containing protein were buffer exchanged back into lysis buffer, concentrated, flash frozen with 10% glycerol, and stored at -80°C.

For purification of GST-tagged TclE, 30-ml cultures were inoculated with 1 ml overnight culture, grown at 37°C until an OD_600_ = 0.6, then IPTG was added and the cells were grown for an additional 20 h at 25°C. The cells were harvested by centrifugation and the cell pellets were frozen at -80°C for at least 30 min. The cells were then thawed and resuspended in 1 ml lysis buffer (50 mM Tris pH 8, 150 mM NaCl, 0.5 mg/ml lysozyme, 2 mM EDTA and one Roche protease inhibitor tablet per 10 ml). Complete lysis was achieved after a 15-min incubation at room temperature, followed by addition of DTT to 1.5 mM. Lysate was processed with several short sonication pulses with a microtip. Insoluble material was centrifuged at 7,000 × g, and the supernatant was combined with 30 µl of glutathione-agarose resin (slurry) at 4°C for 45 min (rotating). The resin was pelleted and the beads were washed with GST buffer three times and the peptide was eluted with 40 µl GST buffer plus 10 mM reduced glutathione. The eluant was either frozen at -80°C for later use, or directly treated with tobacco etch virus (TEV) protease and ZipTipped (using the manufacturer’s instructions).

Activity of Tcl enzymes was tested *in vitro*. 20-μl reactions containing 20 µM GST-TclE, 5 mM DTT, 2 mM ATP, 20 mM MgCl_2_, 1 µM enzymes, and 1 µg TEV protease, were allowed to react for 40 min at RT. Reactions were zip-tipped (using the manufacturer’s instructions) and analyzed by matrix-assisted laser desorption ionization-time of flight (MALDI-TOF) mass spectrometry.

### Orbitrap Liquid Chromatography Mass Spectrometry (LC-MS) for TclE truncation series

GST-TclE leader-truncated samples were co-expressed with TclIJN as described in the Tcl protein expression and purification section. GST-TclE leader truncations were purified using glutathione beads as described above and TclE peptides were removed from the GST tag through TEV protease cleavage. Truncated TclE peptide samples were alkylated to cap any reduced cysteines using chloroacetamide at 20 mM. The peptides were then separated and measured via liquid chromatography-mass spectrometry (LC-MS) on an Easy nLC 1200 in connection with a Thermo Easy-spray source and an Orbitrap Fusion Lumos. Peptides were pre-concentrated with buffer A (3% acetonitrile, 0.1% formic acid) onto a PepMap Neo Trap Cartridge (particle size 5 μm, inner diameter 300 μm, length 5 mm) and separated with an EASY-Spray™ HPLC Column (particle size 2 μm, inner diameter 75 μm, length 25 mm) with increasing buffer B (80% acetonitrile, 0.1% formic acid) gradient:

Samples were eluted using a gradient of 5% B to 22% B over 85 minutes (128 minutes for muscle), 22% to 32% B over 15 minutes (22 minutes for muscle), with a wash of 32% to 95% B over 15 minutes, which was held at 95% B for 15 minutes followed by a wash step consisting of two washes going from 95% B to 2% B over 3 minutes, holding at 2% B for 3 minutes, returning to 95% over 3 minutes and holding for 3 minutes were performed. Sample loading and equilibration were performed using the HPLC’s built in methods. LC-MS only runs were performed using 2400 V in the ion source, scan range of 375-1700 m/z, 30% RF Lens, Quadrupole Isolation, 8 *105 AGC Target and a maximum injection time of 50 ms. The MS-based data-dependent acquisition method was set to a 3 second cycle time. MS1 scans were acquired by the orbitrap at a resolution of 120,000. Precursors with a charge > 1 and < 6 were selected for MS2 fragmentation. MS2 scans of CID precursor fragments were detected with the linear ion trap at a scan rate of 33.333 Da/sec with a Dynamic injection time. CID collisions were set to 30% for 10ms. After 3 selections a 60 second dynamic exclusion window was enabled; isotopes and unassigned charge states were excluded.

### Data Processing for Label-free Quantitation

Raw files were searched against a FASTA data base for the TCL operon (containing I, J, N, and E entries) with the *E. coli* proteome as a contaminant (Uniprot Reference UP000000625) using Peaks Studio analysis. The parent mass error tolerance was set to 10 ppm and the fragment mass error tolerance was set to 0.5 Da. Cysteine carbamidomethylation, thiazoline, and thiazole were set as a variable modification, as well as methionine oxidation and pyro-glu from glutamine were set as variable modifications in the search. Digest mode was set to unspecific, and the peptide length range was set to 6 – 55 amino acids. The false discovery rate (FDR) for peptide matches was set to 1%, and protein ID significance was set to -10log(P-value) ≥ 15. Label-free data was normalized using the TIC option in PEAKS then the total signal for the modified form was compared as a fraction of the total signal for the peptide of interest (**File S1**).

### Modeling and optimization of Tcl protein structures

The sequences of TclI, TclJ, TclN, as well as the protein hetero-dimers such as TclE:TclI, TclI:TclJ and TclI:TclN were submitted online to the AlphaFold2 Google colab (51, 52) for structural predictions. The top-ranked models were selected for each Tcl protein and dimer. Each model underwent optimization using the FastRelax algorithm (53) within PyRosetta. Energy calculations for each model were carried out using the ref2015 score function (54, 55) in PyRosetta. Binding energies were determined by subtracting the energies of the bound and unbound state models. Additionally, to assess the binding modes, we calculated the shape complementarities (56) of the complex models using Python logic and PyRosetta. These models were visualized using PyMOL, and a list of interacting residues (L187R, T188R, Y189R, H194R, I198R, C199R, N201R, I202R, S205R, E206R, F208R, L209R, Y210R, T212R, S213R) was identified for use in mutagenesis experiments.

### Size Exclusion Chromatography of TclI and TclIJN complex

Purified complex was filtered through 0.2 µm cellulose filter (14000 g, 2 min). All the samples were vacuum dried, and then resuspended in SEC buffer (100 mM sodium phosphate with pH 6.8, 0.023% NaN_3_) to make the final concentration to 1 µg/µL. The Agilent 1260 Infinity HPLC System (Agilent, Santa Clara, CA) equipped with quaternary pump, manual injector, thermostatted column compartment, diode array detector (DAD) was used to carry out the analytical size exclusion chromatography. A Yarra-1.8 µm X 150Å, 150 × 4.6 mm HPLC column (00F-4631-E0, Phenomenex, USA) was used for separation of molecules. The Agilent system and column were equilibrated with 100 mM Sodium Phosphate with pH 6.8, 0.023% NaN_3_ at a flow rate of 0.3 mL/min at 25 °C. The molecular weight calibration curve for SEC was obtained by running a protein standard mix containing bovine thyroglobulin (670 kDA), IgA (300 kDa), IgG (150 kDa), ovalbumin (44 kDa), myoglobin (17 kDa) and uridine (0.244 kDa), (AL0-3042, Phenomenex, CA, USA) to relate the molecule’s size to elution volume. The injection volume for the protein standard and the protein samples was 5 µL, and the elution volume was measured using the UV detector. All the data was collected at 280 nm, with a reference wavelength of 600 nm.

### Bioinformatics analysis of E1/YcaO, and Ocin-ThiF-like domains

Protein sequences of TclI and TclJ were submitted to HHpred (36–38) in FASTA format using standard parameters, and PDB70 and TIGRFAMs databases to analyze for the presence of E1/YcaO/Ocin-ThiF-like domains using pairwise comparison of profile HMM (hidden Markov model). The results of this analysis provided a list of known homologs with well-defined domains, and multiple sequence alignments that were used to define each domain on the TclI and TclJ proteins.

## ACKNOWLEDGEMENTS

Research reported in this publication was supported by the National Institutes of Health (NIH) grants 1R15GM132852-01 to JSG and R01AG066874 to JCP. We thank Clarissa Clark and Katherine Brown for their assistance in processing samples for Liquid Chromatography Mass Spectrometry and Size Exclusion Chromatography.

